# High Throughput RNA Sequencing of Germ-Free Mouse Retina Reveals Metabolic Pathways Involved in the Gut-Retina Axis

**DOI:** 10.1101/2020.10.01.318949

**Authors:** Urooba Nadeem, Bingqing Xie, Asadolah Movahedan, Mark D’Souza, Hugo Barba, Nini Deng, Vanessa A. Leone, Eugene Chang, Dinanath Sulakhe, Dimitra Skondra

## Abstract

**Background and aims:** Connections between the gut microbiome and retinal diseases such as age-related macular degeneration (AMD), diabetic retinopathy (DR), retinopathy of prematurity (ROP), and primary open-angle glaucoma (POAG) are recently being established. Communication between the gut microbiome and retina, referred to as the gut-retina axis, has been proposed; however, the biologic pathways and mediators involved in the interactions have not yet been elucidated. Using high-throughput RNA sequencing (RNA-seq) of whole retinas, we compare the retinal transcriptome from germ-free (GF) and specific pathogen-free (SPF) mice to investigate the effects of the gut-microbiome on both retinal gene expression and biologic pathways.

**Methods:** RNA was extracted from whole retinas of GF and SPF mice (four animals per group) and cDNA libraries were created. RNA-seq was performed on NovaSEQ6000 using the paired-end method. After preprocessing the RNA-seq data, gene expression value was calculated by count per million (CPM). The differentially expressed genes (DEGs) were identified with the limma package from Bioconductor on the expression data. Functional enrichment and protein-protein interaction STRING protein-protein association network analyses were created for the differentially expressed genes (DEGs).

**Results:** RNA-sequencing reveals a cohort of 396 DEGs, of which, 173 are upregulated and 223 are downregulated in GF mouse retina. Enrichment analysis reveals that the DEGs are involved in glucocorticoid effects, transcription factor binding, cytoskeletal stability, lipid metabolism, and mitogen-activated protein kinase (MAPK). Multiple biologic pathways, including obesity/metabolic syndrome, longevity, insulin-like growth factor (IGF) signaling pathway, vascular endothelial growth factor (VEGF), hypoxia-inducible factor (HIF)-1 transcription pathway, and 5’ AMP-activated protein kinas**e** (AMPK) signaling pathway are affected in the GF retinas. PPARG1a (PGC1a) gene is involved in 13 of the 35 significantly modulated pathways. Proteins with the greatest number of interactions in the PPI are E1A binding protein P300(EP300), forkhead box O3(FOXO3), and PGC1a.

**Conclusions:** To our knowledge, this is the first study demonstrating the involvement of the gut microbiome in driving the retinal transcriptome, providing evidence for the presence of a gut-retina axis. Future studies are needed to define the precise role of the gut-retina axis in the pathogenesis of retinal diseases.

## Introduction

The gut microbiome is the collective microbiota and their genetic material residing symbiotically in the gut. ^(1)^ Effects of the gut microbiome in diseases of anatomically distant sites, for instance, skeletal muscle, lung, and brain, are well-recognized. ^(2,3).^ However, its role in ocular conditions, especially retinal diseases is only recently being recognized. ^(4,5)^ Communication between the gut microbiome and retina, also referred to as the gut-retina axis has been proposed; but to date, the biologic mediators and pathways affected in the retina by the gut microbiome are not known.

The concept of the gut microbiome affecting anatomically distant organs, especially the regulation at an immune-privileged site, like the eye, is very perplexing. ^(6)^ In the past few years, data for the gut microbiome’s involvement in retinal diseases is accumulating. ^(7,8)^ Autoimmune uveitis was the first ocular disease linked with an altered gut microbiome. ^(9)^ Early microbiome studies show germ-free (GF) mice completely devoid of microbiota, have a modified retinal lipidome. ^(10)^ Huihui Chen *et al.* showed that GF mice relative to mice with an intact microbiome are protected from glaucoma by a T-cell mediated effect. ^(11)^ Alteration of the microbiome by diet (also known as gut dysbiosis) illustrates that high-sugar and fat diets exacerbate worsened features of dry age-related macular degeneration (dAMD) and neovascular AMD, respectively. ^(4,12,13)^ On the other hand, it is shown that restricting the gut microbiome by fasting prevents diabetic retinopathy (DR) and prolongs survival in mice. ^(14)^ Human studies have determined that the gut microbiome is compositionally and functionally different in wet AMD, ^(14)^ primary open-angle glaucoma (POAG), ^(15)^ retinal artery occlusion (RAO) ^(17)^ and retinopathy of prematurity (ROP) ^(18)^ in contrast to healthy subjects. Gut microbiota modulation may serve as a novel strategy for the prevention and treatment of retinal diseases.

A recent study shows that remodeling the gut-microbiome has a favorable influence on retinal morphology and decreases age-related retinal ganglion cell (RGC) loss. ^(19)^ Another investigation suggests that administration of probiotics may modulate clinical signs of autoimmunity in the eye. ^(20)^ Andriessen et al show the gut microbiota influences the development of neovascular lesions associated with AMD and modifying microbiota can decrease systemic and local chorioretinal inflammation, subsequently decreasing pathological choroidal neovascularization. ^(12)^ Despite this knowledge, the gut microbiome’s precise role in retinal disease pathogenesis remains unknown; nonetheless, the functional differences between health and disease states support the presence of a gut-retina axis.

Multiple animal models have been used to study the microbiome, using antibiotics and diet to modulate the microbiome are the most common methods for these studies. ^(4, 12)^ Nevertheless, these models do not allow for assessing changes caused by antibiotics and diet, as opposed to the effects caused by the microbiome alone. ^(21)^ 16S rRNA gene sequencing can be used to compare taxonomic differences between gut microbes of patients and healthy controls. ^(22)^ All these techniques provide significant primary evidence for linking the gut-microbiome to the retinal diseases; but, to move beyond the correlation phase of experiments; preclinical mouse models are essential. ^(21,23)^ The GF mouse model is considered the gold-standard for proof-of-concept microbiome investigations and has been used in milestone discoveries to determine causal links between the microbiome and disease. ^(24,25,26)^ Presently, no studies use the GF mouse model to describe the gut microbiome’s role in the gut-retina axis.

Here, we profiled the retinal transcriptome to identify differentially expressed genes (DEG) between GF and specific pathogen-free (SPF) mice retina; the latter serving as a control group. Advances in high throughput RNA-sequencing and computational bioinformatics analysis have evolved as powerful methods to yield detailed information on novel genes and pathways involved in different conditions. ^(27)^ The study design is illustrated in Figure 1. To our knowledge, no microbiome studies have illustrated alterations in the retinal transcriptome utilizing RNA-seq technology. We identified DEGs between the two groups revealing dysfunctional molecular functions, biological processes (BPs), cell components, and pathways. Furthermore, protein-protein associations of the DEGs are also explored.

**Figure1.**
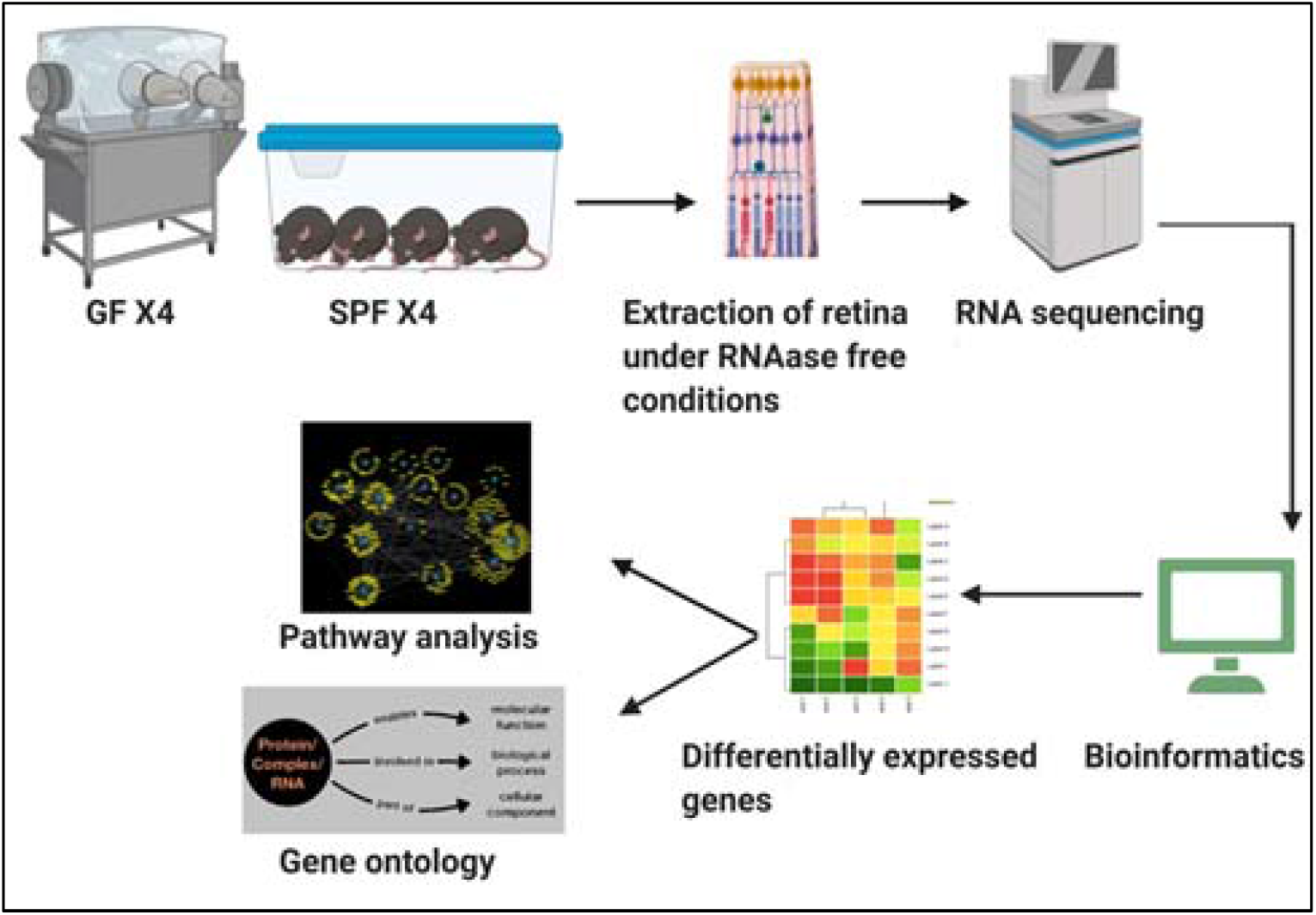
Schematic diagram illustrating the experiment methodology. Image made using Biorender.

Changes in gene expression by modulating the gut-microbiome have already been established in the GI tract, ^(28)^ skeletal muscle, ^(29)^ and the liver. ^(30)^ We show that similarly the gut microbiome can also modulate gene expression in the retina. Our study provides a global insight into the retinal genes and pathways influenced by the gut microbiome. Future investigations are needed to elucidate the function of the microbiome in specific retinal diseases, which can aid in developments in alternate or adjunct prevention strategies and therapies for medically refractory and incurable retinal diseases.

## Materials and Methods

### Animals

All experiments were performed per the Association for Research in Vision and Ophthalmology (ARVO) statement for the use of animals in Ophthalmic and Vision Research and were institutionally approved. Adult male mice C57Bl/6 mice were utilized for this study. The GF mice were bred in the Gnotobiotic Research Animal Facility (GRAF) at the University of Chicago (U of C). SPF mice were purchased from the Jackson Laboratories and bred at the Animal Resources Center (ARC) at the U of C SPF animal vivarium. SPF mice were kept in sealed cages in a ventilated rack system.

Mice were fed normal rodent chow. Food and water were provided *ad libitum* to both groups and they are exposed to 12:12 light: dark cycle. The environmental parameters of temperature and humidity were kept within the range required by The Guide for the Care and Use of Laboratory Animals, Eighth Edition. (31) At 15 weeks of age, the mice were sacrificed via CO2 asphyxiation followed by cervical dislocation. All measures were taken to minimize stress and pain throughout the procedure. Sample processing for this experiment was immediately started on ice as an extra cautionary measure to prevent biologic/ genetic material contamination and degradation.

### Sterility Monitoring

To ensure sterility in GF mice, they were housed, bred, and maintained in a sterile flexible film isolator at the U of C GRAF. Diets for GF mice were autoclaved (250 F, 30 minutes). Weekly fecal pellets were collected for sterility monitoring via microbial cultures, as previously described. (32) Positive cultures were not identified in any of the GF mice used in this study.

### RNA extraction

After the mice were sacrificed, the eyes were enucleated and carefully dissected, and the retinas were extracted on ice using a sterile technique. All surfaces, tools, tubes, and equipment were treated with RNase decontamination solution (RNaseZAP, Thermofisher Scientific Waltham, MA). The retinas were immediately stored in RNAlater (Qiagen, Germantown, MD) in RNase free Eppendorf tubes and stored at −80C until RNA extraction. For the mRNA extraction, the RNeasy (Qiagen, Germantown, MD) was used. Each sample was analyzed using Nanodrop (NanoDrop 2000cc, Thermo Scientific, Waltham, MA) to determine total RNA concentration before sequencing.

### RNA sequencing

Eight samples of purified RNA (4 from each group) were used for this analysis. Before sequencing, the RNA quality was assessed via the Bioanalyzer, and the samples were confirmed to have the required RNA integrity numbers (RIN) concentration (both done at U of C Genomics core). The cDNA library of each sample was prepared using Tru-Seq RNA Sample Prep Kits (Illumina, San Diego, CA) for 100 bp paired-end reads, according to the manufacturer’s instructions. Each of the eight cDNA libraries were indexed for multiplexing. These indexed libraries were sequenced on NovaSEQ6000 (Illumina, San Diego, CA) using PE100bp. Data were recorded in the FASTQ format and then imported in R for bioinformatics analysis.

### Statistical Analysis

The secondary analysis of sequence data was performed on Globus Genomics,^(33)^ an enhanced, cloud-based analytical platform that provides access to different versions of Next-Generation Sequence analysis tools and workflow capabilities. Tools such as STAR^(34)^, featureCounts,^(35)^, and Limma were run from within the Globus Genomics platform. We used STAR (version 2.4.2a, Stanford University, CA) aligner default parameters to align the RNA-seq reads to the reference mouse genome (GRCm38) for all eight samples. The raw gene expression count matrix was then generated by featureCounts (version subread-1.4.6-p1). ^(35)^ The gene annotation was obtained from the Gencode vM23. ^(37)^ STAR default parameter for the maximum mismatches is 10 which is optimized based on mammalian genomes and recent RNA-seq data.

We obtained a matrix with 55213 genes and eight samples as the final output. The data was imported to R for downstream analysis. The raw gene count matrix for all samples was transformed into log-CPM (count per million) values. Genes without sufficiently large counts were filtered using EdgeR Bioconductor. ^(38)^ 20,287 genes were selected for the downstream analysis. We used Limma Voom ^(39)^ normalization to standardize the gene expression matrix, where the raw library sizes were scaled using TMM (trimmed mean of M values). Multidimensional scale plots were generated on top 6 leading FC dimensions which showed the similarity between samples.

The differential expression analysis was performed on contrast group SPF against GF using Limma. (40) Significant DEGs with p-value < 0.01 were extracted for further downstream analysis. The enrichment analysis in the Toppgene suite took both the upregulated and downregulated DEGs in GF and extracted the over-represented gene ontology functional classification (molecular functions, biological processes, and cellular component). ^(41)^ The significance of the association between the datasets and bio functions were measured using a ratio of the number of genes from the dataset that map to the pathway divided by the total number of genes in that pathway. This enrichment analysis was based on mouse-to-human orthologs. A list of all DEGs and their p-values is available in Supplementary Table 1.

Proteins are known to work together to exert a specific function. Therefore, we mapped the identified DEGs into the STRING (Search Tool for the Retrieval of Interacting Genes) database (http://string-db.org/) version 11.0, which contains a wealth of validated and text mined protein-protein associations among the proteins, to further explore the relationships of these DEGs from protein level. ^(42)^ We used the DEGs (p-value < 0.01) to identify protein-protein associations for both the upregulated and downregulated genes in GF mice.

## Results

### Gut microbiome modifies global retinal gene expression

To compare the retinal transcriptome profile of SPF vs.GF mice, we performed high throughput RNA-seq analysis of mouse retinas from the two groups. To obtain precise results, four separate retinas were obtained from each sample group (biological replicates). After the correction of the raw data to remove background noise, 20,287 genes were selected for differential gene analysis. Given the false discovery rate (FDR) were less than 0.51 in the differential gene expression, the significant differentially expressed genes were selected based on a stringent p-value cutoff 0.01. DEGs were labeled as the genes with a p-value of <0.01. A comparison between the two groups revealed that 396 DEGs were identified, 173 upregulated and 223 downregulated genes in the GF mice group. Using the criteria of log fold-change (log_2_FC)>1, we performed an unsupervised analysis of these DEGs. This revealed 60 genes; 40 upregulated and 20 downregulated in the GF mice retina. Heatmap are plotted to show the hierarchical clustering of these genes (Figure 2). The volcano plot demonstrates a distinct retinal gene expression in retinas from SPF vs GF mice (Figure 2). Our data supports that there are substantial differences in the global gene expression in GF and SPF retina; hence, supporting the presence of gut-retina axis. Detailed statistics of the DEGs are available in Supplementary Table 1.

**Figure2.**
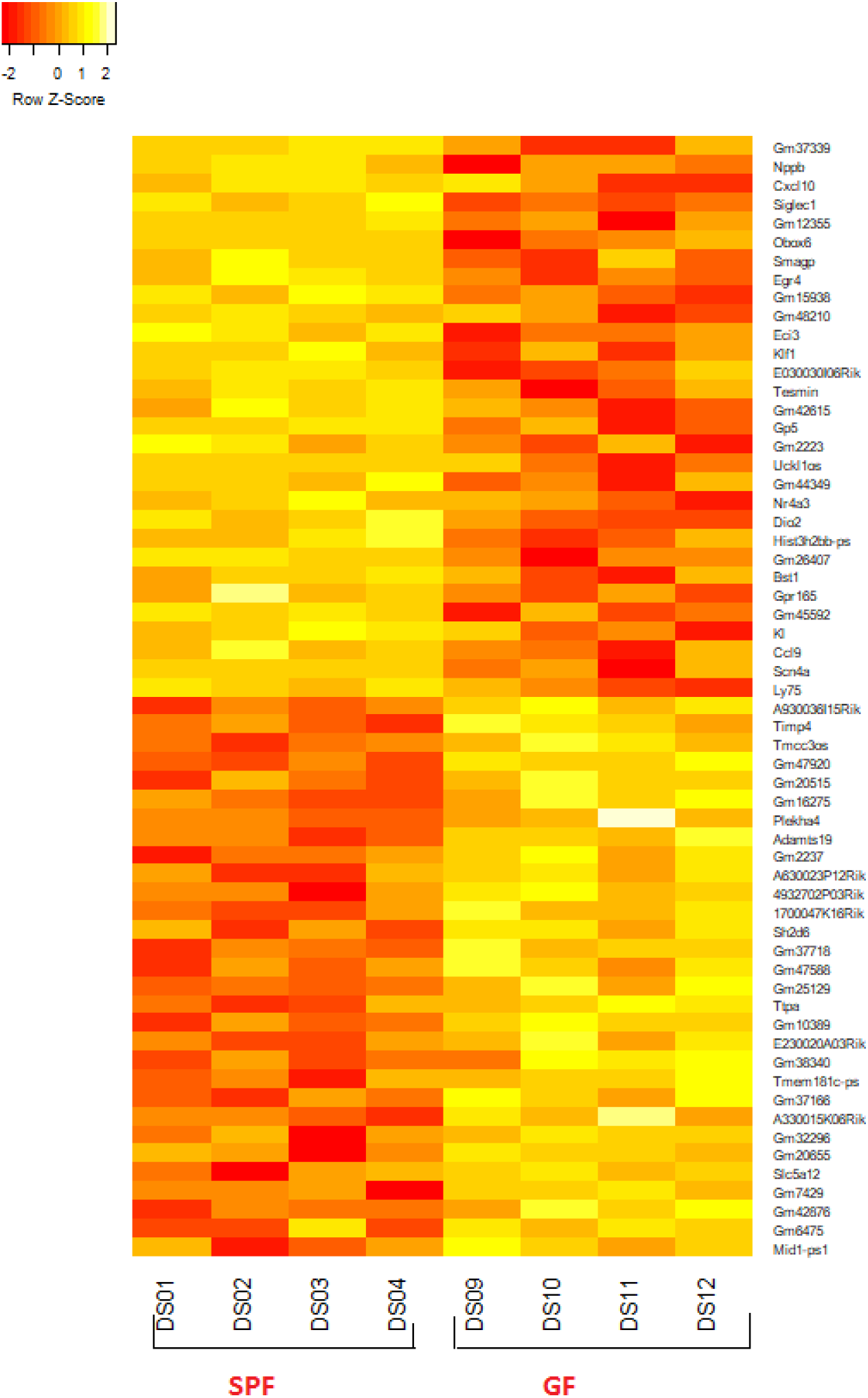
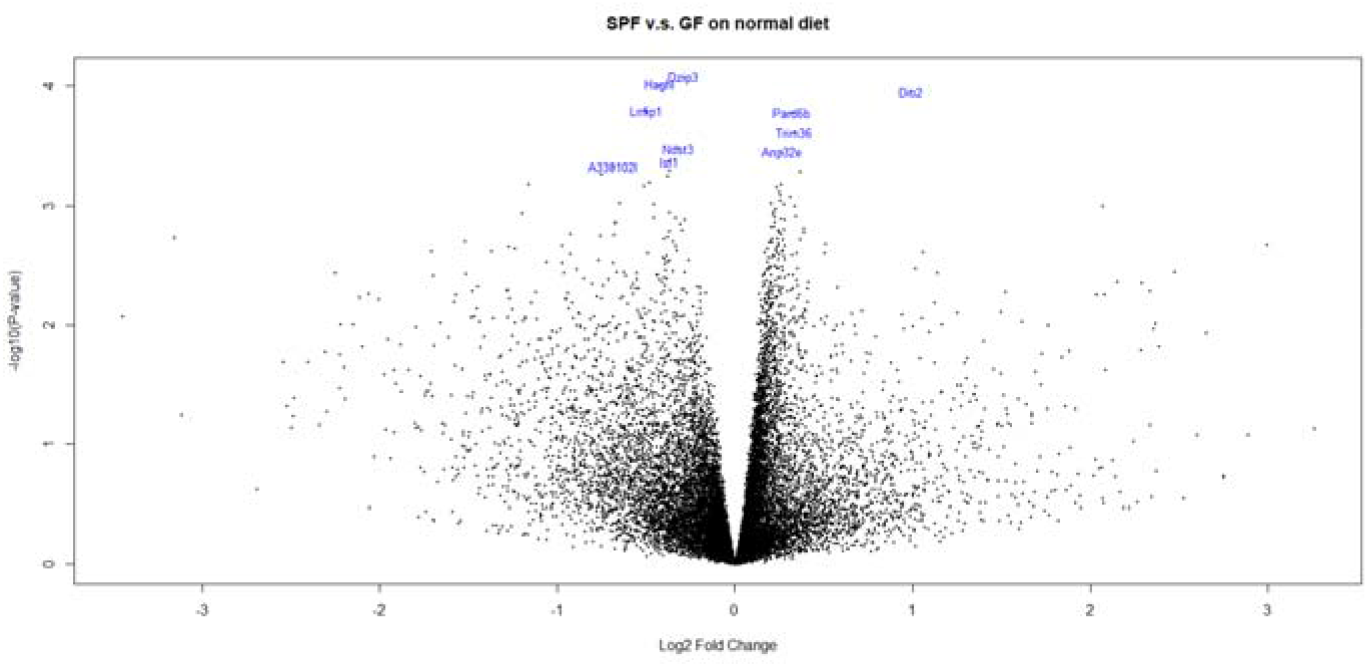
Comparing the gene expression profiles of the GF and SPF mice. (A). Hierarchical clustering of retinal genes of GF vs SPF mice demonstrated by the heatmap. Red and yellow indicate upregulated and downregulated genes in the GF mice retina, respectively. DEGs with log2FC > 1 (more than 2-fold difference) are included in this heatmap. (B). Volcano plot showing DEGs in GF vs SPF mice shows a distinctive gene expression profile of two groups with minimal overlap. X-axis represents log2 FC and the Y-axis represents −log10 (P-values). (DS01, DS02, DS03, DS04-SPF and DS09, D10, DS11, DS12-GF)

### Enrichment Analysis reveals Gut Microbiome Regulates Multiple Key Biological Pathways in Mouse Retina

The enrichment analysis for both Gene Ontology and pathways was performed using Toppgene ^(41)^. The genes were mapped from mouse ortholog. In total, 302 genes out of 396 genes were mapped by the ortholog. The enrichment analysis identifies statistical significantly (FDR<0.05) over-represented gene ontology functional categories (Figure 3) and pathways based on DEG (Figure 4). The analysis revealed that the DEGs are involved in glucocorticoid receptor binding (GO:0035259), steroid hormone effects (GO:0071383), transcription factor binding (GO:0008134), cytoskeletal stability (GO:0007010), lipid metabolism (GO:0071396) and mitogen-activated protein kinase (MAPK) activity (GO:0017017).

**Figure 3A, B, C.**
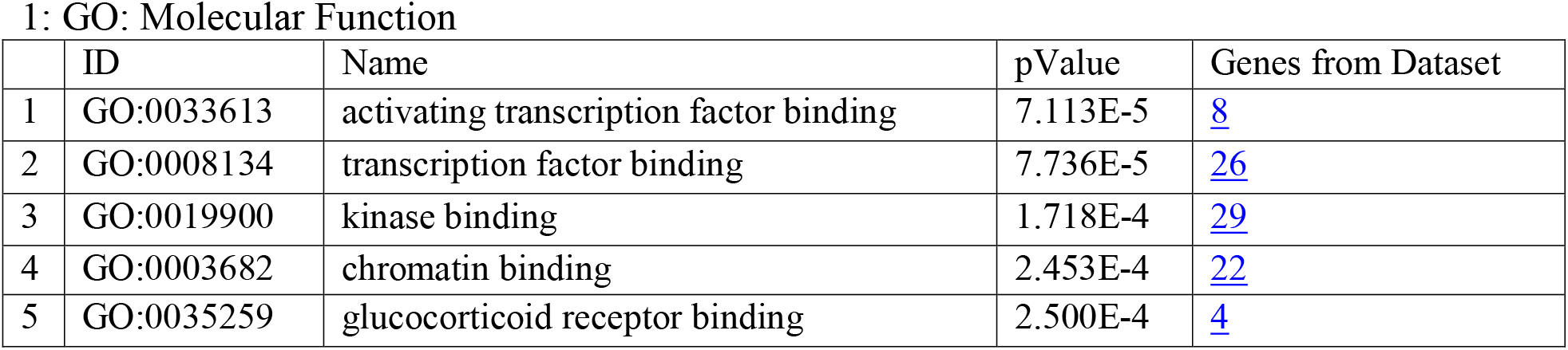
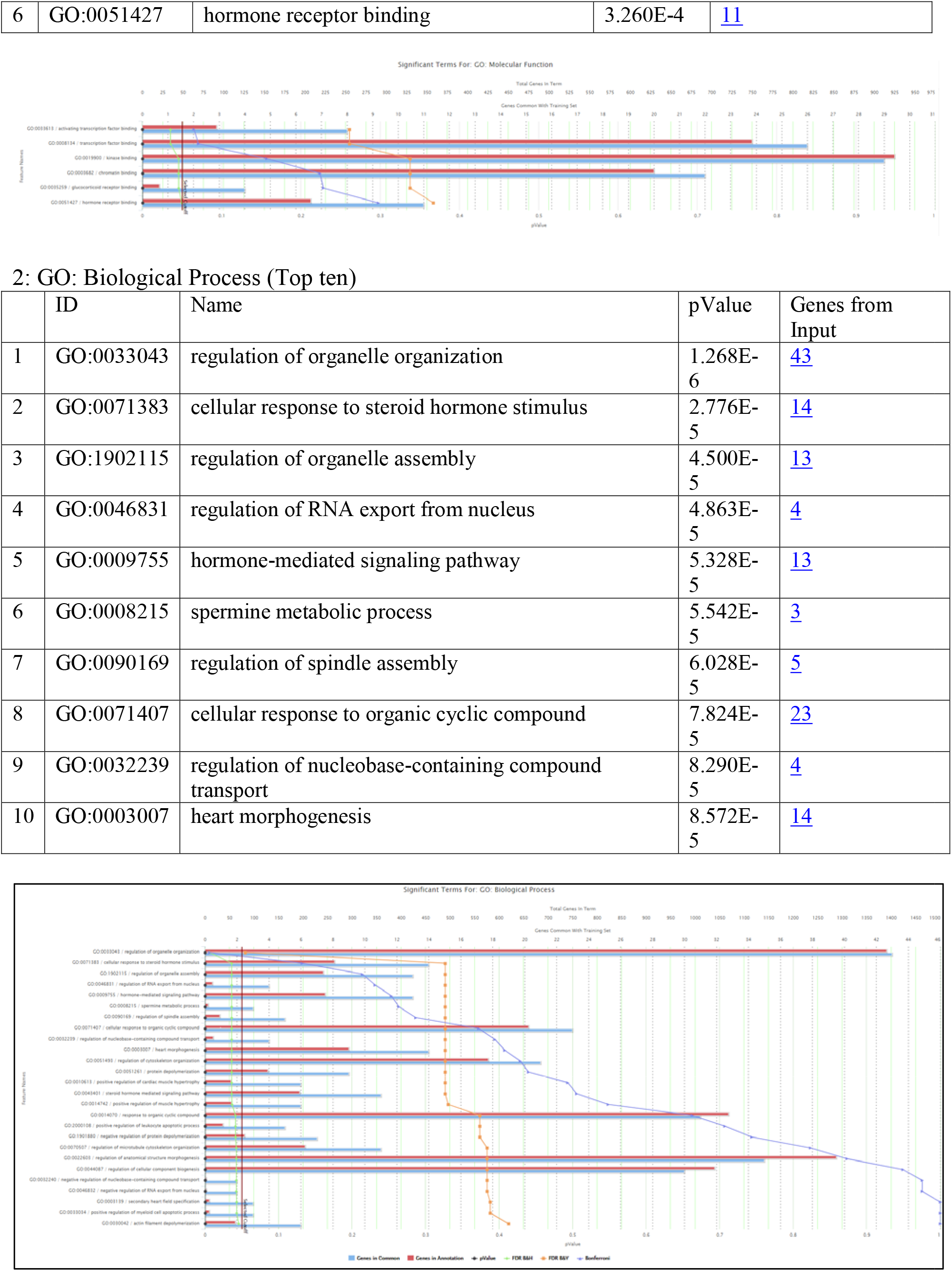
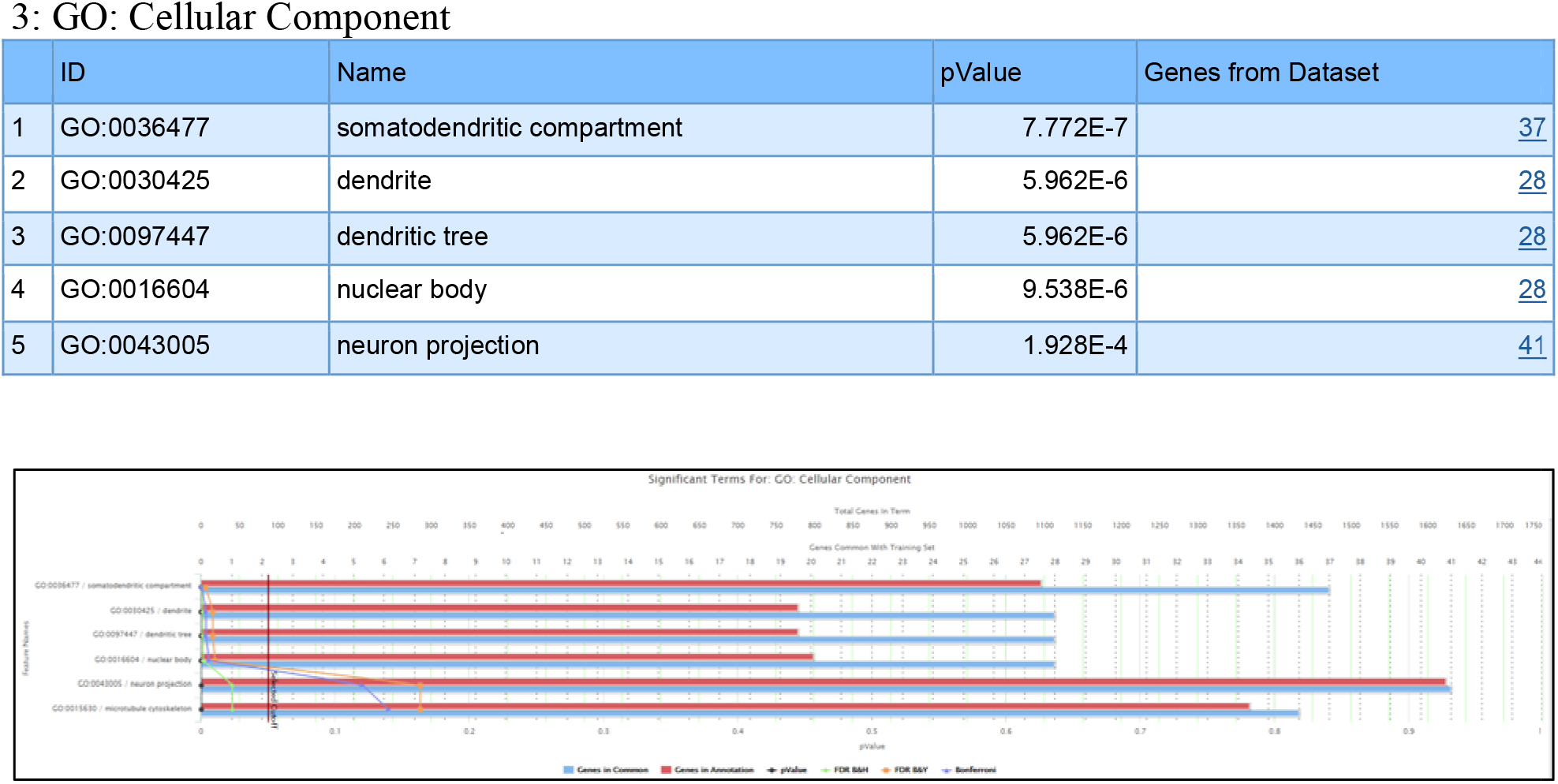
Table for top 10, molecular functions, biologic processes and cell component revealed to be different in the GF mouse retina by enrichment analysis in the DEGs. Corresponding bar-charts show the genes affected in our dataset compared to the total number of genes involved in the condition. The figure is made using Toppgene.

**Figure 4:**
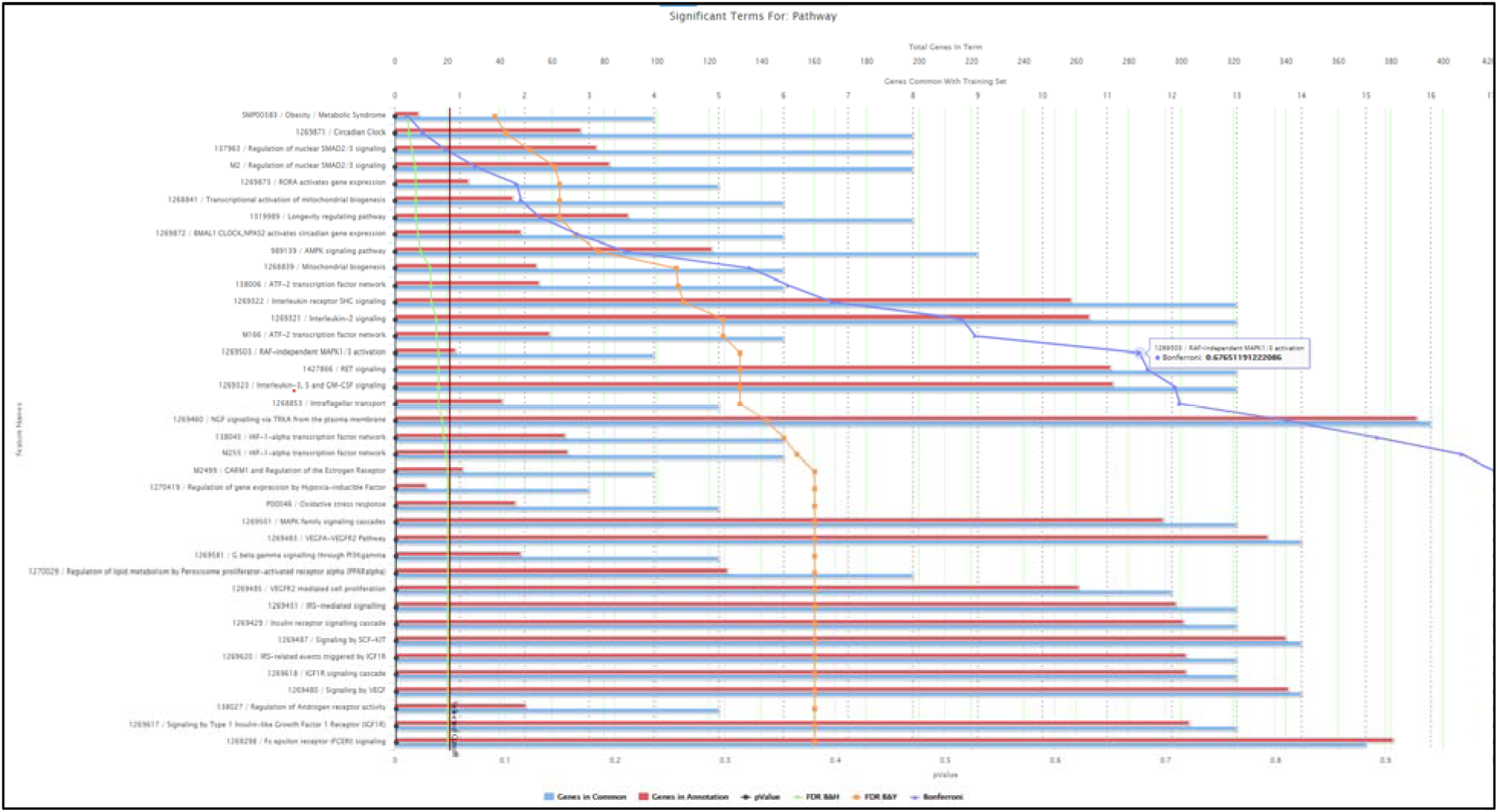
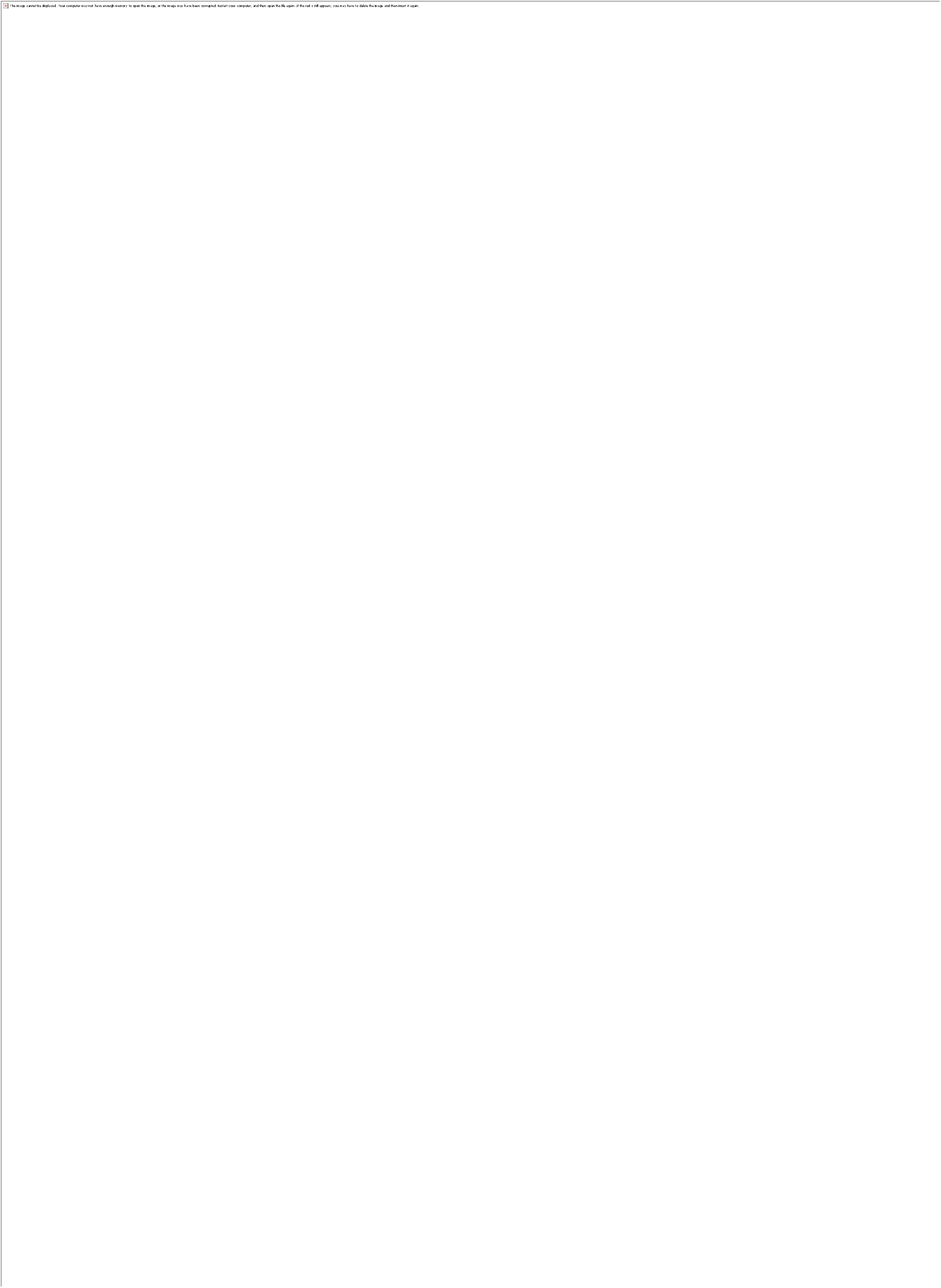
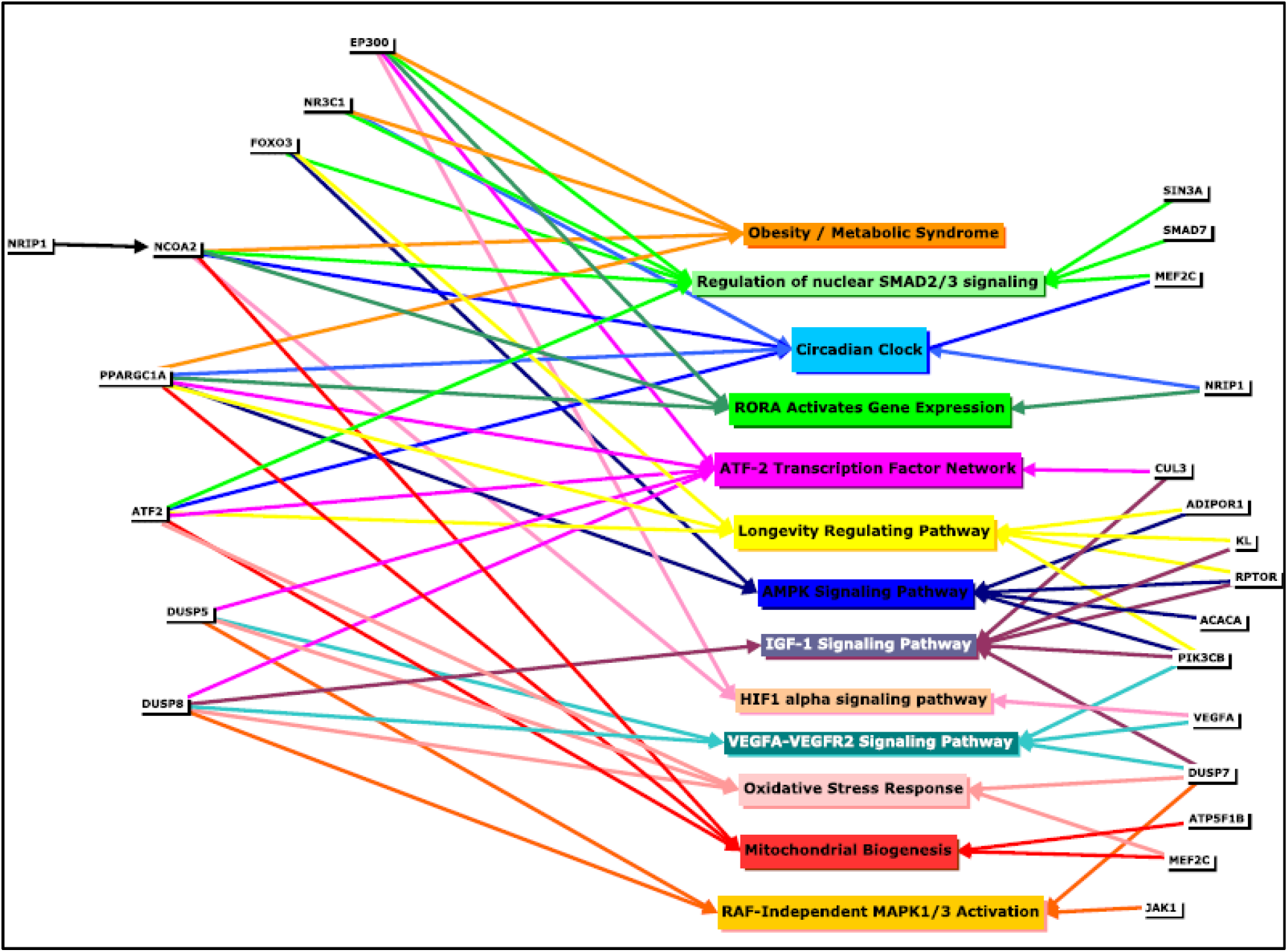
The pathways downregulated by the DEGs revealed by enrichment analysis. Corresponding bar-charts showing the genes affected in our dataset compared to the total number of genes involved in the GF mice. The figures are made using Toppgene. **Figure 4b**. Select pathways and their corresponding involved genes revealed by enrichment analysis. The downregulated pathway is in the middle of the figure. Genes involved in multiple pathways are represented on both the left and right of the figure.

The absence of the gut microbiome influenced 35 biologic pathways in the GF retina (Figure 4). Since Toppgene uses data from multiple different databases to determine the relevant pathways from the list of the genes, the pathways that appear more than once by the original enrichment analysis are removed. These pathways were involved in obesity/metabolic syndrome, adenosine monophosphate-activated protein kinase (AMPK), nuclear SMAD 2/3 signaling, mitochondrial biogenesis, lipid metabolism, longevity, both vascular endothelial growth factor (VEGF) mediated cell proliferation, hypoxia-inducible factor (HIF)-1 transcription, IGF-1and oxidative stress response, all of which are already established to be crucial in the pathogenesis of numerous retinal diseases. However, novel pathways with no known function in retinal diseases, but demonstrated to be regulated by the gut-microbiome, for example, intraflagellar transport is also shown to be affected by the gut microbiome. Our data shows that the DEGs in the modified pathways have considerable overlap (Figure 4). PPARGC1A (now referred to as PGC-1α) is involved in 13 of the 35 pathways and seems to be especially important.

### Protein-Protein associations

Using the STRING database with both the upregulated and downregulated DEGs in the GF retina groups, protein-protein association analysis was done. In the protein-protein association network, a node represents a protein and the degree of a node represents the number of the interplayed pairs with other proteins of this specific node. The proteins which are the predominant nodes with a high degree of connections to other proteins (hub nodes) include histone acetyltransferase p300 (EP300), PGC1a, VEGF, nuclear receptor coactivator 2 (NCOA2) and forkhead box O3 (FOXO3) in the analysis conducted on the DEGs downregulated in the GF mice. In this network, there are total 308 proteins and 379 edges where the average node degree is 2.46. The node degree for selected the hub proteins is higher than 10 and the major hub EP300 has 23 direct neighbors in this network. PGC1a, FOXO3, Vegfa and NCOA2 have 12, 14, 10 and 10 direct neighbors, respectively.

The pathway analysis of the all the proteins perturbed in the GF retina compared to the SPF retina shows that they are involved in are AMPK signaling and longevity regulating pathways (Supplementary table 4).

## Discussion

In this study, we used high throughput RNA sequencing to compare the entire retinal transcriptome of whole retinas from GF and SPF mice. Our results show that retinal gene expression is altered by the gut-microbiome the presence of gut microbiotas. 396 DEGs (p-value <0.01) are identified between the two groups. Functional categories the DEGs belong to include glucocorticoid receptor binding, steroid hormone response, lipid metabolism, and transcription factor binding. Additional genes involved in cytoskeletal stability and regulation of T-helper-17 (Th17) differentiation are also affected. Analysis of the DEGs shows pathways affected by the gut-microbiome are obesity/metabolic syndrome, MAPK, mitochondrial biogenesis, longevity regulation, AMPK, IGF-1, VEGF, HIF-1 transcription factors, and oxidative stress (Figure 4). Molecules proved shown to be regulated by the gut-microbiome in the gastrointestinal tract like tissue metallopeptidase inhibitor 4 (43) and mucin 2 (44) are also affected in the GF mice retina. Hub nodes in the downregulated PPI network include PGC1a, EP300, and FOXO3; all of which are involved in the pathogenesis of retinal diseases. Notably, the pathways predicted to be downregulated in GF retina analysis by STRING analysis are AMPK and longevity pathways, further suggesting that these pathways are indeed influenced in the retina by gut microbial status.

We hypothesized that the gut microbiome regulates retinal function by modifying retinal gene expression. To test the hypothesis, we use GF mice which are considered the gold standard for proof-of-concept questions for microbiome studies. As discussed above, GF mice are important for demonstrating correlative relationships. (21,22,25) They have proved invaluable for establishing causal links between the microbiome and diverse diseases such as inflammatory bowel disease, cancers, autoimmune conditions, and neurologic conditions. (45,46) Recent evidence highlights the importance of the gut microbiota in modulating response to different cancer immunotherapy drugs. (47) Methods for studying the microbiome include microbiota inhibition using antibiotics, diet alteration to cause gut dysbiosis (changes microbial composition), and the introduction of microbial strains directly into animals that have conventional microbiota. (48) Modulating the microbiome through diet and/or antibiotics is relatively easier, cheaper, and does not require specialized facilities compared to the GF mice. Nonetheless, using antibiotics has limitations, including off-target drug effects, only partial ablation of microbes, possible selection of resistant strains, and direct influence of the drug on the genetic and metabolic machinery of host tissue (48). Dietary manipulation results in alteration of the microbiota; however, a high-fat diet may cause obesity regardless of the composition of gut microbiota in mice. (49) Therefore, in addition to the indirect dietary influence caused by dysbiosis, it is likely that there is a direct impact of the dietary constituents on the host tissue. Introducing microbial strains by fecal transplant has revealed to be beneficial in ocular surface conditions, e.g. it shows an improved goblet cell density, decreased corneal barrier disruption, and dacryoadenitis in animal models of Sjogren’s syndrome. However, only very limited studies comparable investigations have been attempted for retinal diseases. Host-microbe interactions can be studied more accurately and precisely without the background noise from a preexisting microbiome in the GF mice. (21,22) Hence, we chose GF mice to identify genes and pathways involved in the gut-retina axis. To our knowledge, this is the first report using GF mice to outline the transcriptomic changes in the retina caused by changes in the gut microbiome.

As mentioned earlier, there are compositional and functional differences in the enteric microbiome in AMD, POAG, RAO, and ROP patients. Investigators are trying to comprehend how different species of microbiota contribute to genomic changes in the host. Numerous theories have been proposed; the most accepted are the downstream impact from the metabolites produced from the microbiota. Taylor *et al*. show that modifying the gut-microbiome by a high-glycemic-index diet leads to retinal damage; they use metabolomics to compare the severity of the retinal damage with multiple microbial metabolites. (4) Other studies show the protective nature of the gut microbiota metabolites in the retina; exogenous short-chain fatty acids (SCFA) decreases autoimmune uveitis,(50) ursodeoxycholic acid (UDCA) ameliorates DR (51), and tauroursodeoxycholic acid (TUDCA) protects against cell death in retinal degeneration, (52,53)retinal detachment,(54) and protects neural retina in the diabetic mouse model. (55) Further credence is provided to this theory by the presence of SCFA transporters, sodium◻coupled monocarboxylate transporter 1 and 2 (SMCT1 and SMCT2), and receptor G protein-coupled receptor 109A (GPR109A) in the retina. (56) Our data shows that SMCT2 (SLC5A12) gene is upregulated in GF mice (Supplementary Table 1). Another recent study shows that modifying the gut microbiome by feeding mice *L.paracasei KW3110* demonstrates favorable changes in retinal function and morphology on the aging retina. (19) These studies lay the groundwork for the connection between the enteric microbiome and the retina; however, these investigations did not reveal if the microbiota or its metabolites may alter the retinal gene expression. RNA-seq is a revolutionary technique that can generate very precise results to study the entire transcriptome of the tissue. In our study, we used high throughput RNA-seq to evaluate the retinal transcriptome of GF mice retina. We generated a detailed view of the genes and pathways affected in the retina by the gut microbiome but further studies are required regarding the mechanism by which gut microbiota regulates the retinal gene expression as seen in our studies.

The importance of DEGs in retinal diseases including AMD, POAG, ROP, and DR (Figure 6) is well-recognized. The most influenced gene in our data is the transcriptional coactivator PGC-1α, which is affected in 12 of the 16 altered pathways (figure 4B). This gene is involved in numerous retinal diseases. Zhang *et. al* show that AMPK/PGC-1α pathway is dysfunctional in AMD patients. They show that alteration of this pathway leads to increased intracellular accumulation of lipids and cellular waste, decreased mitochondrial biogenesis and turnover, and subsequent mitochondrial dysfunction. (57) Our data shows both PGC-1α and AMPK pathways are differentially expressed in the GF mice retina, indicating that the gut microbiome is involved in the regulation of this pathway in the retina. Additional enrichment analysis reveals genes involved in downstream activities of AMPK/PGC-1α including lipid metabolism and mitochondrial biogenesis are also influenced (Figure 5). AMD is a heterogeneous disease with a very unpredictable course; if the differences in gut-microbiome cause AMPK/PGC-1α dysregulation and explain this heterogeneity in different patients remains to be seen. PGC-1α is also the major transcriptional coactivator for steroid hormone receptors and its nuclear receptors. Expectedly, glucocorticoid receptor binding, and cellular effects of steroids are among the top ten dysfunctional molecular functions and biologic processes affected in GF mice (Figure 3). We show for the first time that and PGC-1α-steroid pathways are both modified in the retina through the gut-microbiome. However, further studies of protein expression and hormone activity are needed to determine the microbiome’s influence on PGC-1α protein in models of specific retinal diseases.

**Figure 5.**
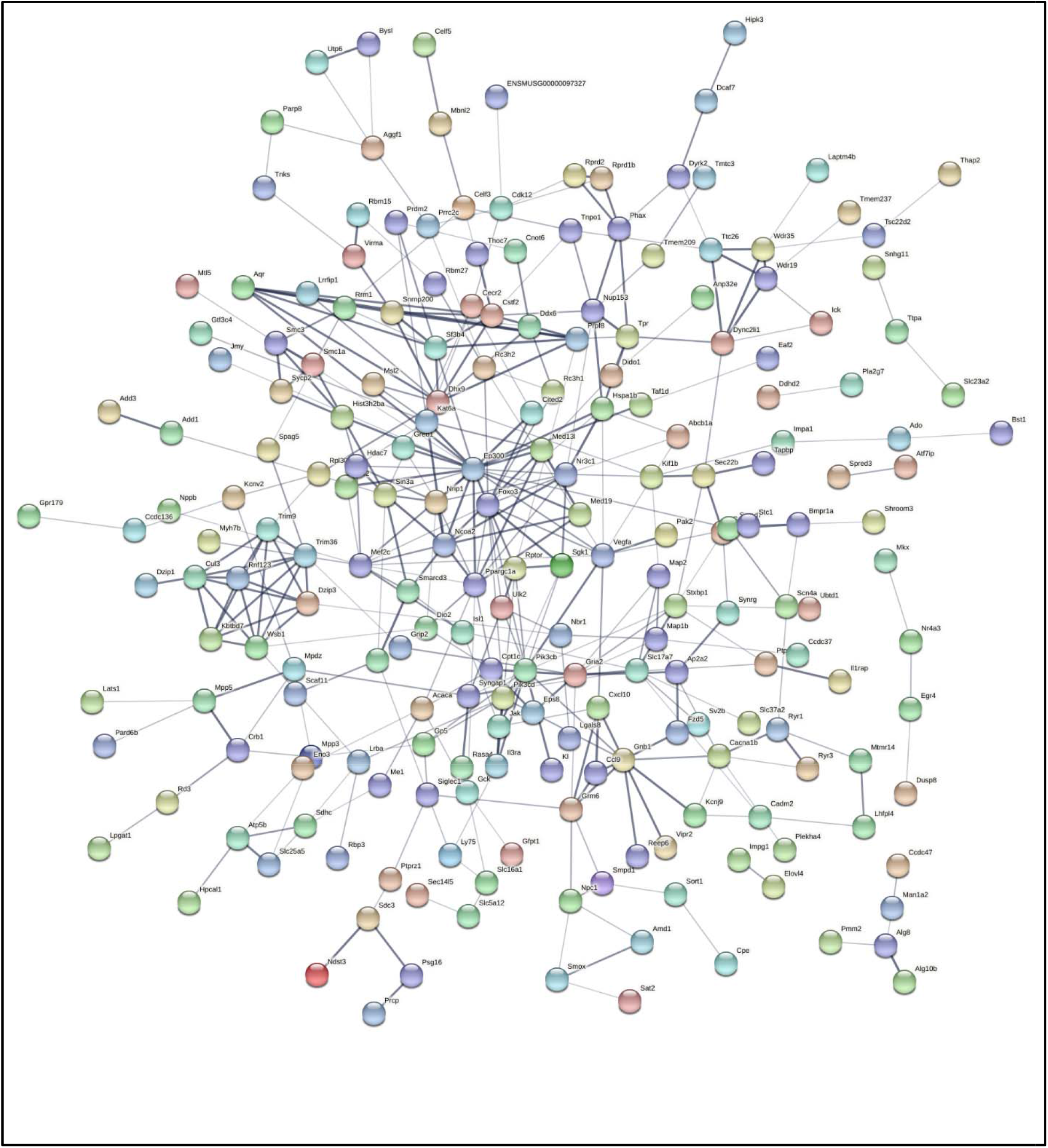
String networks of protein-protein interaction (PPI) generated using DEGs (p-value < 0.01). The figure is generated using *STRING* version 11.0. EP300, PGC1a, FOXO3, NCO2 and VEGF are the major hub nodes.

**Figure 6.**
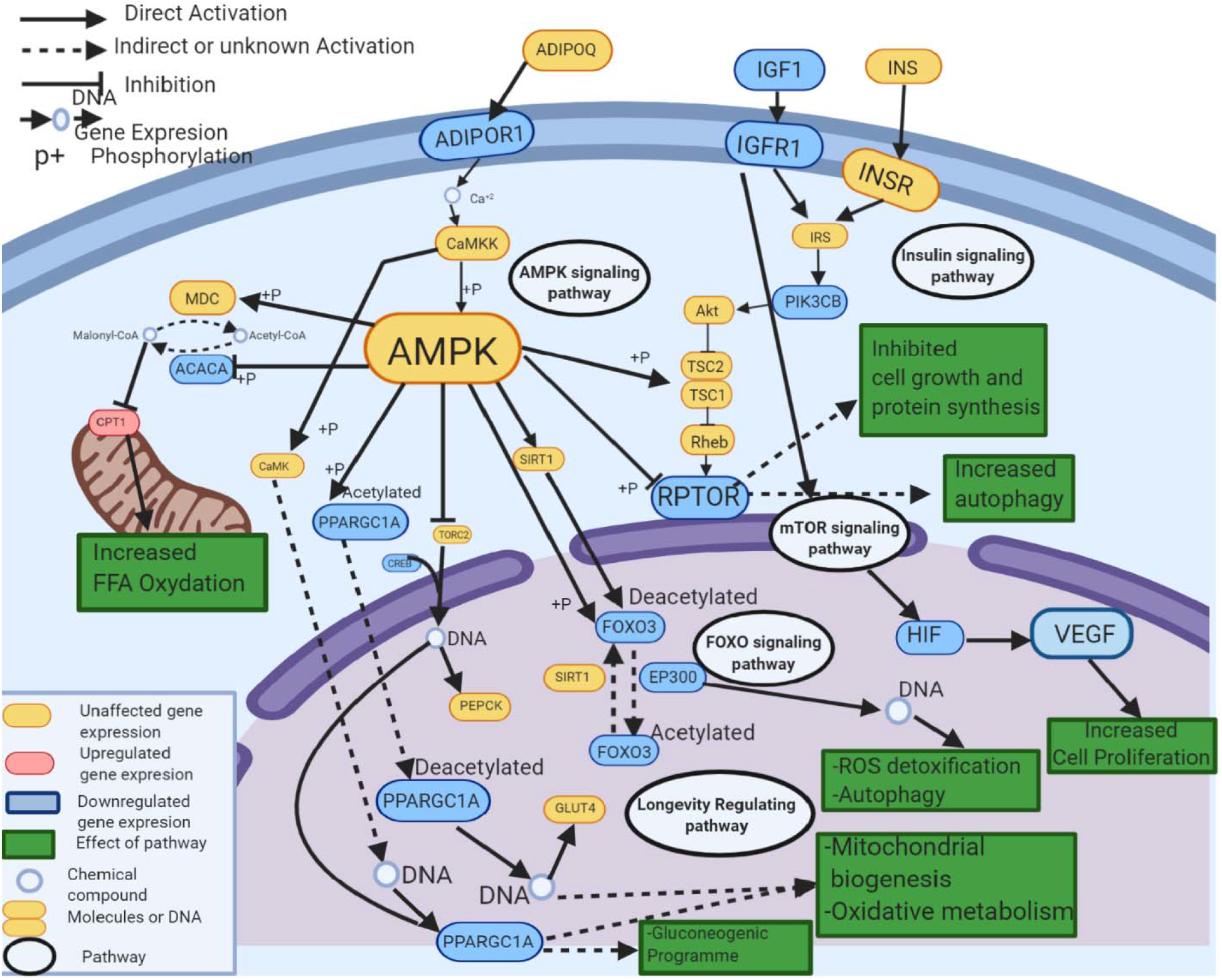
AMPK and the genes and pathways perturbed by the gut-microbiome in GF mice. Image made using Biorender.

AMPK can also function through genes other than the PGC-1A (Figure 6). One of these principal genes is RPTOR (Regulatory Associated Protein of MTOR Complex 1) which is the crucial regulator of the mTOR pathway. The RPTOR gene is downregulated in the GF mice retina (Supplementary Table 1). The mTOR is essential for multiple metabolic pathways and cycles including protein activation, lipid anabolism, and inhibiting autophagy. Overactivity of mTOR is believed to retinal diseases including, AMD,^(57)^ glaucoma,^(58)^ and DR.^(59)^ The mTOR signaling is activated by hormones and growth factors, e.g. insulin and IGF-1. These anabolic factors activate the phosphatidyl inositol-3-phosphate kinase (PI3-k) pathway and protein kinase B (PKB/Akt) leading to mTOR phosphorylation and activation (Figure 6). ^(60,61)^ mTOR activation leads to subsequent HIF 1 nuclear translocation ^(62)^ and increased protein translation from the HIF-1 gene.^(63)^ Increased HIF finally culminates in the increased production of VEGF. Today, targeting VEGF through intravitreal injections is the mainstay to delay the risk of progression in angiogenesis-driven eye diseases, like wet AMD and DR.^(64, 65,66,67,68, 69)^ Our data indicate that the entire AMPK-mTOR-IGF-1-HIF-1-VEGF pathway is affected by the gut microbiome in the retina (Table 4, Figure 6). Hence, highlighting a possible mechanism of how the gut microbiome affects the retinal metabolic pathways. Therefore, suggesting that modifying the gut microbiome can have a profound influence on multiple pathways simultaneously.

In the context of retinal diseases, this finding can help elucidates major concepts in disease pathogenesis. The retina is considered as a projection of the CNS due to the common embryologic origin via neuroectoderm and anterior neural tube. Claud et. al also demonstrated the IGF-1 pathway’s association with neuron and oligodendrocyte development and its involvement in the cross-talk between the gut microbiota and brain using the same GF mouse model as ours. ^(70)^ This suggests there is an overlap between the pathways that are affected in the brain by the gut microbiome and the gut-retina axis, which based on similar embryology is conceivable. Furthermore, very preterm newborns who develop severe ROP are more likely to have evidence of brain damage, including cerebral palsy, developmental delay, lower scores on measures of verbal and performance IQ. ^(71)^ ROP, like AMD, is a heterogeneous disease at the opposite end of the age spectrum; nevertheless, not all high-risk preterm infants develop ROP or acquire the same severity of the disease. Recently, members of our team demonstrated that there is a significant enrichment of Enterobacteriaceae in preterm infants who develop type 1 ROP, hence demonstrating that gut microbiome differences at the taxonomic level. ^(18)^ Low levels of circulating serum IGF-1 are strongly associated with ROP ^(72)^ and we show that the IGF-1 pathway is modulated by the gut-microbiome, at the transcriptomic level. Thus, providing a possible mechanistic explanation of how the gut microbiome is involved in the pathogenesis of severe ROP development. But further studies are needed to delineate these precise associations between the microbiome, brain, and retinal development and growth.

The other important thing concept is the modulation of the HIF1 transcription network as well as both VEGF mediated cell proliferation and signaling in the retina by the gut microbiome. HIF-1 activation via hypoxia is the most important activator of VEGF protein production. Today, the role of the HIF-1-VEGF axis in the angiogenesis of retinal neovascular and degenerative diseases like neovascular AMD, ROP, and DR is almost indisputable. ^(73,74)^ Gut dysbiosis via a high-fat diet leads to an increase in the size of neovascular AMD lesions and a concurrent increase in VEGF-A levels in the choroid. ^(12)^ Also, increased VEGF-A expression has been shown to exacerbate dry AMD features.^(75)^ AMD is the leading cause of blindness in individuals over 60 years of age and there is neither a complete cure nor treatment to prevent AMD. ^(76)^ The initial insult leading to over-activation of VEGF pathways in AMD has not yet been discovered but is believed to be multifactorial-genetic predisposition, aging and diet have all play a role, but the mechanistic pathways linking these predisposing factors toward diseases progression are yet to fully understood.^(77)^ Based on our data, we hypothesize that the gut microbiome may have an important contribution in the pathogenesis of distinct neovascular and degenerative retinal diseases via regulation of HIF-1 and VEGF pathways, in addition to other pathways of the gut-retina axis we describe in this study. Can the gut-microbiome be the important missing piece of the puzzle of explaining the pathogenesis of complex retinal diseases. An exciting aspect of these findings and their possible implications in disease pathogenesis, as well as future applications, is that the gut microbiome is easily modifiable. Theoretically, a targeted gut microbiome modification can modulate multiple pathways simultaneously, unlike most pharmacologic agents that target one pathway or molecule at one time. This can provide a new treatment strategy to tackle multifactorial diseases like AMD. At this time, the field of microbiome studies in the retina field is in its infancy and more studies are needed about which specific microbial profiles and strains, and metabolites involved in microbial-host interactions may be beneficial for improving long-term outcomes in patients with retinal neovascular and degenerative diseases.

Another downregulated pathway in the GF mouse retina is the Smad2/3 signaling pathway. The Smad signaling pathway is synergistic with the MAPK. MAPK phosphorylates multiple transcription factors, including c-Jun and ATF-2 and Smad proteins. Phosphorylation by MAPK pathways results in subsequent activation of their transcriptional activity. Predictably, each of these transcription factor genes and MAPK are downregulated in the GF retina. There is crosstalk between MAPK signaling and several different pathways, such as VEGF and oxidative stress.^(78)^ Oxidative stress via reactive oxygen species (ROS) serves as an important signal that activates MAPK signaling, which is also dysregulated in the GF eyes. MAPK activation has been implicated in the pathogenesis of several retinal diseases, including AMD and DR. However, despite this, several ocular MAPK inhibitors have failed to produce desired outcomes in retinal diseases; in fact, MAPK drugs for tumors may have serious ocular side effects.^(79)^ Given our data, the regulation of MAPK via the gut-microbiome may serve as a probable target for intervention in retinal diseases.

A recent study also shows melatonin inhibits inflammation and apoptosis in animals with DR via the MAPK pathway.^(80)^ A separate investigation reveals that the gut microbiota is sensitive to melatonin and expresses endogenous circadian rhythmicity.^(81)^ Our study supports that several retinal genes of the circadian rhythm clock are affected by the gut microbiome. The mammalian retina contains a circadian clock system independent from the central clock located in the suprachiasmatic nucleus (SCN) of the brain.^(82)^ Studies show that circadian clock dysregulation is also associated with obesity. Dysregulation of the circadian clock also regulates the WNT/β-catenin pathway, malfunction of which is also connected to exudative wet AMD. ^(83)^ The gut microbiome regulates circadian rhythm in the brain;^(84)^ but, we provide the first evidence that the gut microbiome can also affect the circadian circuit in the retina.

Another important set of genes that are affected in GF mice retina is involved in longevity. Currently, only two genetic factors have been consistently found to contribute to longevity; apolipoprotein E (Apo-E) and FOXO3. ^(85)^ Predictably, the analysis of PPI of the downregulated genes in the GF mice retina also highlights FOXO3 as a hub node involved with several other proteins. Aging is the most significant factor contributing to the pathogenesis of retinal degenerative diseases, like AMD. FOXO3 transcription factor is involved in cell survival and apoptosis. PI3K/Akt inhibitors via FOXO3 have demonstrated some efficacy in ameliorating neovascularization in mice models. Gut dysbiosis induced by a high-fat diet leads to the inactivation of FOXO3 in the intestine. Our data indicate that the gut microbiome can affect the gene expression of FOXO3 in the mouse retina, but more studies are needed to determine the protein expression and activity of these genes in the retina.

Gut-microbiome is well-known to cause downstream sequelae by mediating both innate and adaptive elements of the immune system.^(86)^ GF mice demonstrate an increase in T helper cell type 2 (Th2) compared to T helper cell type 1 (Th-1) cells and a decrease in T helper 17 cells (Th-17), T regulatory cells (Tregs), and IL-12 formation.^(87)^ Similarly, the Th17 differentiation pathway in GF mice retina is affected through RC3H1, RC3H2, SMAD7 genes. Both AMD and POAG patients have increased the Th1/ Th17 response. ^(88,89)^ Several genes of the cytokine-cytokine receptor interaction are also affected by the gut-microbiome in the retina. Membrane cofactor protein (CD46), a regulator of the alternative pathway of the complement system, is also downregulated in the GF mice retina. CD46 decrease can lead to hyperactivation of the complement system and it has been established that there is a central role of inflammatory pathogenesis in AMD. Retinal pigment epithelial (RPE) cells, the main cell implicated in the AMD pathogenesis, lose their CD46 expression in geographic atrophy even before the appearance of morphologic changes.^(89)^ Cd46−/− knockout mice are used as an experimental animal model who spontaneously develops a dry-type AMD-like phenotype.^(90)^ Our experiment reveals that both innate and adaptive immunity pathways are disrupted in the GF mice retina. Despite most agents targeting complement system seem promising in theory and in vitro, their efficacy is not satisfactory when evaluated in clinical trials. To date, there are no successful clinical trials in providing a drug that effectively controls the inflammation in the retina. ^(91)^ Since the gut-microbiome affects both innate and adaptive immune pathways it is possible that discovering the precise organisms that modulate the inflammatory pathways can be exploited for future therapeutic benefits.

Several long non-coding RNAs (lncRNA) are differentially expressed in the GF and SPF mice retina (Supplementary Table 1). Long non-coding RNAs (lncRNAs) are increasingly recognized to control cellular functions and regulate transcriptional and translational processes of the neighboring genes. Gut microbiota critically regulates the expression of lncRNAs not only locally in the intestine but also remotely in other metabolic organs, like brown adipose tissue and muscle.^(92)^ The exact role of lncRNA in either the retinal disease is unclear at the moment; however, they have an important function in the retinal ganglion cell differentiation. Our study indicates that the gut microbiome could affect retinal functions via regulation of retinal lnc-RNAs expression and possible downstream effects on neighboring genes as seen in other organs.

Nonetheless, the GF mouse studies have some limitations since host physiological parameters are altered in these mice. The lack of microbiota produces systemic changes in these animals, including the inability to efficiently digest nutrients and having an under-developed immune system. Other variations include lengthier intestinal epithelial turnover, lower nutritional requirements, and less body fat despite increased consumption. Moreover, our data is based on RNA-seq alone, and no studies of protein expression or activity were performed. RNA-seq studies typically provide the basis for further experiments using specific disease animal models and subsequent human diseases.

## Conclusion

In this study, we explored the retinal transcriptome using high-throughput RNA-seq to investigate the effect of the gut microbiome on the entire retinal transcriptome of the GF mice to determine the genes and pathways involved in the gut-retina axis.

Our data provide strong evidence for the regulation of important pathways including AMPK, VEGF, HIF-1, IGF-1, MAPK, longevity, oxidative stress, and mitochondrial biogenesis by the gut-microbiome in the retina. Hence, we propose that regulation of biologic retinal pathways via the gut microbiome may be crucial in retinal diseases, especially AMD, DR, ROP, and other degenerative retinal diseases. The gut-microbiome caused modulation in gene expression has been demonstrated in other organs; but, this is the first report providing solid evidence of the existence of the gut-retina axis and the important biological pathways involved. Even though our data indicate that the gut microbiome is a powerful regulator of retinal gene expression, further analyses are needed to elucidate how retina protein expression and cellular functions are modified by the gut-microbiome and their correlation with retinal disease pathogenesis.

## Acknowledgments

We thank University of Chicago Core Genomic team, Peter Fabier and Mikayla Marchuk, for helping with RNA sequencing. Gnotobiotic Research Animal Facility (GRAF) staff (Betty Theriault and Melanie Spedale) and Donna Arvans for care of the germ-free animals and Chang lab members Candace Cham, Xiaorong Zhu, Jason Koval and Amal Kambal.

